# Discovering and prioritizing candidate resistance genes against soybean pests by integrating GWAS and gene coexpression networks

**DOI:** 10.1101/2022.05.26.493550

**Authors:** Fabricio Almeida-Silva, Thiago M. Venancio

## Abstract

Soybean is one of the most important legume crops worldwide. Soybean pests have considerable impact on crop yield. Here, we integrated publicly available genome-wide association studies and transcriptomic data to prioritize candidate resistance genes against the insects *Aphis glycines* and *Spodoptera litura*, and the nematode *Heterodera glycines*. We identified 171, 7, and 228 high-confidence candidate resistance genes against *A. glycines, S. litura*, and *H. glycines*, respectively. We found some overlap of candidate genes between insect species, but not between insects and *H. glycines*. Although 15% of the prioritized candidate genes encode proteins of unknown function, the vast majority of the candidates are related to plant immunity processes, such as transcriptional regulation, signaling, oxidative stress, recognition, and physical defense. Based on the number of resistance alleles, we selected the ten most promising accessions against each pest species in the soybean USDA germplasm. The most resistant accessions do not reach the maximum theoretical resistance potential, indicating that they might be further improved to increase resistance in breeding programs or through genetic engineering. Finally, the coexpression networks generated here are available in a user-friendly web application (https://soypestgcn.venanciogroup.uenf.br/) and an R/Shiny package (https://github.com/almeidasilvaf/SoyPestGCN) that serve as a public resource to explore soybean-pest interactions at the transcriptional level.

## 1 Introduction

Soybean (*Glycine max* (L.) Merr.) is the world’s main legume crop, with a primary impact in human and animal nutrition, and in industrial applications. However, soybean fields are significantly affected by pests (insects and nematodes) that lead to dramatic yield losses. The major pests in soybean fields are the soybean aphid (*Aphis glycines* Matsumura) and the soybean cyst nematode (*Heterodera glycines* Ichinohe), which are responsible for annual losses of US$4 billion and US$4.5 billion in the US, respectively (Bandara et al., 2020; Koenning and Wrather, 2010). In Brazil, the world’s leading soybean producer, insect pests reduce yield by 7.7%, which corresponds to an economic loss of US$ 21.78 billion (April 2022 exchange rate) (Oliveira et al., 2014).

Over the past few years, many genome-wide associations studies (GWAS) have been performed to identify single-nucleotide polymorphisms (SNPs) associated with soybean resistance to insect and nematode pests (Liu et al., 2019; Hanson et al., 2018; Zhao et al., 2017; Bao et al., 2014; Natukunda et al., 2019). However, as GWAS typically cannot accurately pinpoint causative genes, multi-omics data integration has helped predict high-confidence candidate genes associated with traits of interest (Baxter, 2020; Michno et al., 2020). Recently, we identified high-confidence candidate genes against fungal diseases using cageminer, a graph-based algorithm recently developed by our group to integrate GWAS and transcriptomic data to prioritize candidate genes (Almeida-Silva and Venancio, 2021b, 2022a). Thus, we hypothesize that our algorithm can also reveal high-confidence candidate genes that can be used to engineer soybean lines with increased resistance to pests.

Here, we integrated multiple publicly available RNA-seq and GWAS datasets to identify high-confidence candidate genes associated with resistance to pests. We found a high overlap of resistance genes between insects, but not between insects and nematodes, suggesting that these classes trigger different defense responses. The candidate resistance genes against each species are involved in several immunity-related processes, such as transcriptional regulation, signaling, oxidative stress, recognition, and phytohormone metabolism. Strikingly, 15% of the candidates encode proteins of unknown function, revealing a novel catalog of potential resistance genes. Finally, we highlighted the ten most resistant accessions against each pest species in the USDA germplasm, uncovering important information for breeding programs and genetic engineering initiatives. The coexpression networks resulting from this work were also made available as a web application (https://soypestgcn.venanciogroup.uenf.br/) and R/Shiny package (https://github.com/almeidasilvaf/SoyPestGCN).

## 2 Materials and Methods

### 2.1 Curation of resistance-associated SNPs and pan-genome data

SNPs with significant association to resistance against soybean pests were manually curated from published GWAS data (Table 1; Supplementary Table S1). SNPs that were present in the SoySNP50k database were identified with their standard nomenclature, and the VCF file for the SoySNP50k was downloaded from Soybase (Brown et al., 2020). Additionally, a matrix of gene presence/absence variation (PAV) in the pan-genome of cultivated soybeans (*n* = 204 genomes from 24 countries and 5 continents) (Torkamaneh et al., 2021) was also used.

**Table 1.**
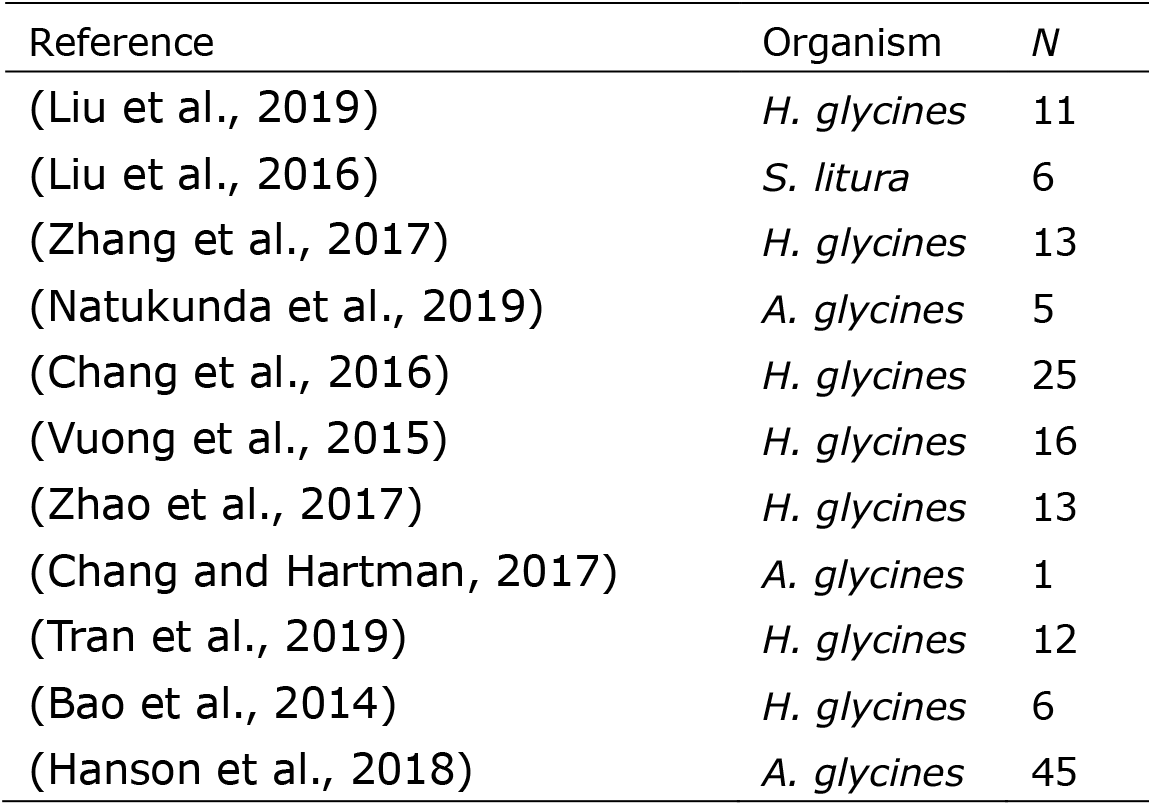
GWAS included in this work. *N*, number of significant resistance-related SNPs in each study.

| Reference | Organism | *N* |
| --- | --- | --- |
| (Liu et al., 2019) | *H. glycines* | 11 |
| (Liu et al., 2016) | *S. litura* | 6 |
| (Zhang et al., 2017) | *H. glycines* | 13 |
| (Natukunda et al., 2019) | *A. glycines* | 5 |
| (Chang et al., 2016) | *H. glycines* | 25 |
| (Vuong et al., 2015) | *H. glycines* | 16 |
| (Zhao et al., 2017) | *H. glycines* | 13 |
| (Chang and Hartman, 2017) | *A. glycines* | 1 |
| (Tran et al., 2019) | *H. glycines* | 12 |
| (Bao et al., 2014) | *H. glycines* | 6 |
| (Hanson et al., 2018) | *A. glycines* | 45 |

### 2.2 Prediction of variant effects on genes

Variant effect prediction was performed with the function *predictCoding()* from the R package VariantAnnotation (Obenchain et al., 2014). Genome sequences and transcript coordinates were downloaded from PLAZA 4.0 (Van Bel et al., 2018). Reference and alternate alleles were manually extracted from each GWAS publication. Variants with no information on reference and alternate alleles in the original publication were discarded from this analysis.

### 2.3 Transcriptome data and selection of guide genes

Gene expression estimates in transcripts per million mapped reads (TPM, Kallisto estimation) were retrieved from the Soybean Expression Atlas (Machado et al., 2020). Additional RNA-seq samples comprising soybean tissues infested with pests were retrieved from NCBI’s SRA in a recent publication from our group (Almeida-Silva and Venancio, 2022b). We filtered the GWAS and transcriptome datasets to keep only insect and nematode species that were represented by both data sources. We selected a total of 102 and 36 RNA-seq samples from soybean tissues infested with insects and nematodes, respectively (Supplementary Table S2). Finally, genes with median expression values lower than 5 were excluded to attenuate noise, resulting in a 15684 *x* 102 gene expression matrix for insects, and a 10240 × 36 matrix for nematodes. Guide genes were obtained from the Supplementary Data in (Almeida-Silva and Venancio, 2021b).

### 2.4 Candidate gene mining and functional analyses

Gene expression data were adjusted for confounding artifacts and quantile normalized with the R package BioNERO (Almeida-Silva and Venancio, 2022a). An unsigned coexpression network was inferred with BioNERO using Pearson’s r as correlation. Candidate genes were identified and prioritized using the R package cageminer (Almeida-Silva and Venancio, 2021a) with default parameters. Module enrichment analyses were performed with BioNERO, using functional annotations from the PLAZA 4.0 database (Van Bel et al., 2018). Finally, prioritized candidates were given scores and ranks using the function *score_genes()* from cageminer.

### 2.5 Selection of candidate resistant accessions from the USDA germplasm

The VCF file with genotypic information for all accessions in the USDA germplasm was downloaded from Soybase (Brown et al., 2020). For each locus *i*, scores S_i_ 0, 1, or 2 were given based on the number of SNPs associated with resistance. Total resistance scores for each accession were calculated as the sum of scores S_i_ for all *n* loci as follows:

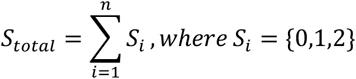

Total resistance scores were ranked from highest to lowest, and ranks were used to select the most resistant accessions. The resistance potential of the best accessions was calculated as a ratio of the attributed scores to the theoretical maximum score (*2n*, which corresponds to all loci having scores 2).

## 3 Results and discussion

### 3.1 Data summary and genomic distribution of SNPs

After removing pest species that were not represented by both soybean RNA-seq and GWAS, our list of target species included the insects *A. glycines* (soybean aphid) and *S. litura* (armyworm caterpillar), and the soybean cyst nematode *H. glycines* (Figure 1A). SNPs associated with resistance to all pest species were located in gene-rich regions of the soybean genome (Figure 1B), and their distributions were biased towards particular chromosomes (Figure 1C). Resistance SNPs against *A. glycines* were mostly located on chromosome 13, and resistance SNPs against *H. glycines* were mostly located on chromosomes 18, 8 and 7 (Figure 1C). Resistance SNPs against *S. litura* only occurred on chromosomes 12, 7, 6 and 5, but it is important to mention the small number of resistance SNPs against this species.

**Figure 1.**
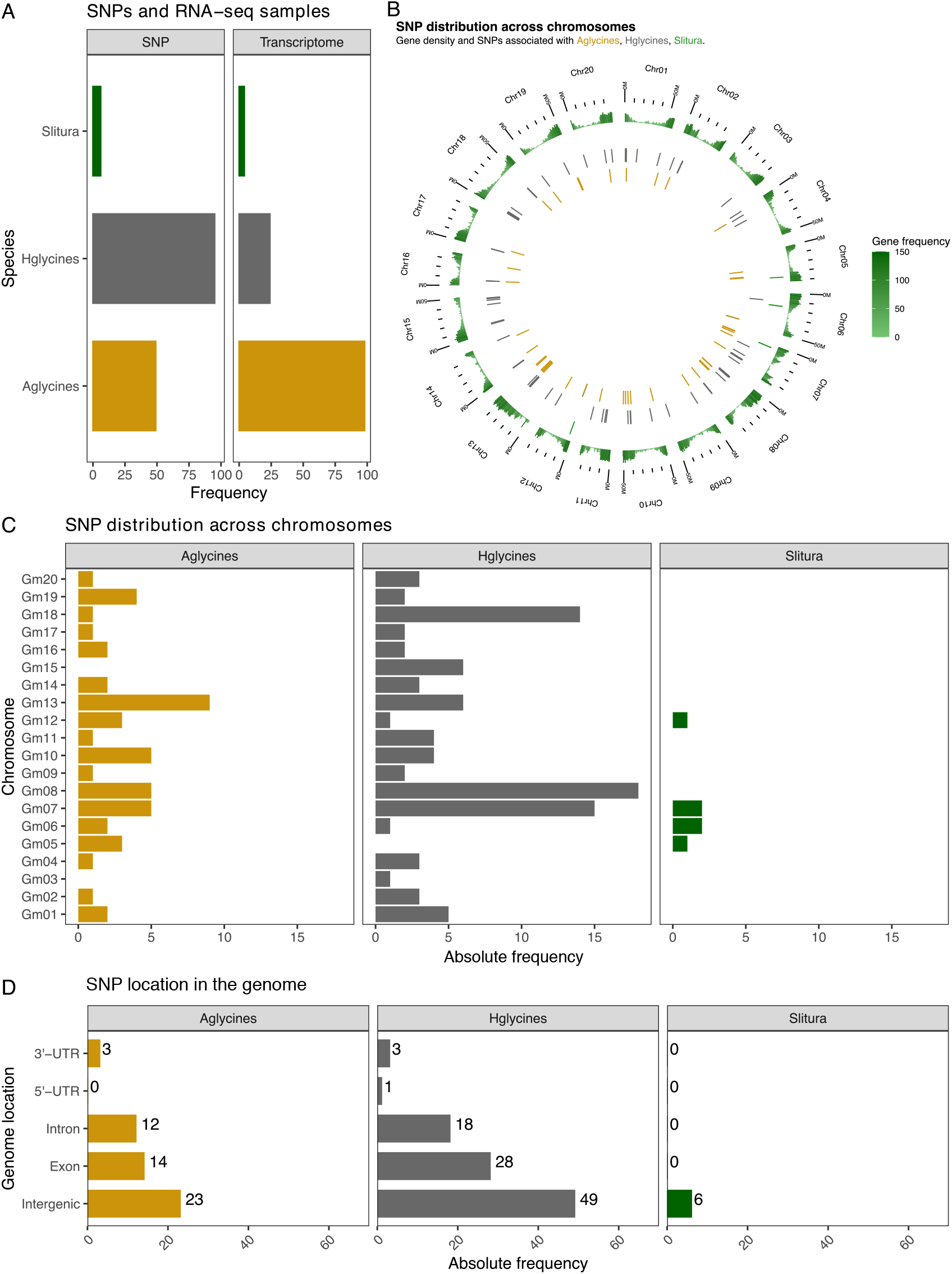
Data summary and genomic distribution of SNPs. A. Frequency of SNPs and RNA-seq samples included in this study. B. Genomic coordinates of resistance SNPs against each pest species. The outer track represents gene density, whereas inner tracks represent the SNP positions for each species. C. SNP distribution across chromosomes. Overall, there is an uneven distribution of SNPs across chromosomes. D. Genomic location of SNPs. Most SNPs are located in intergenic regions.

Interestingly, although most resistance SNPs against all species were located in intergenic regions, a considerable fraction of them was located in exons, except for *S. litura* (Figure 1D). This is a dramatic difference from what we observed in our previous study on fungi resistance-related genes, where almost all SNPs were located in intergenic regions (Almeida-Silva and Venancio, 2021b). Thus, we predicted SNP effects on coding sequences to better understand the functional consequences of these SNPs. From all resistance SNPs against *A. glycines* in coding regions, 31% (*n*=5) led to nonsynonymous substitutions, while 69% (*n*=11) led to synonymous substitutions (Supplementary Table S3). Conversely, from all resistance SNPs against *H. glycines* in coding regions, nonsynonymous substitutions prevailed as expected (63%, *n*=10), followed by synonymous (31%, *n*=5) and nonsense substitutions (6%, *n*=1).

Additionally, we explored the distribution of SNPs in introns to understand their functional impact. SNPs in splice sites (*i.e*., ±2 nucleotides relative to the exon-intron junction), as they can influence exon configuration and alternative splicing (Woolfe et al., 2010). None of the SNPs in introns were located in splice sites, indicating that they do not affect splicing patterns directly. As introns can contain regulatory elements, such as enhancers and silencers (Cooper, 2010), these SNPs might contribute to increased resistance by affecting intron-associated transcriptional regulation.

### 3.2 High candidate gene overlap between insects, but not between insects and nematodes

Using defense-related genes as guides, we identified 171, 7, and 228 high-confidence genes against *A. glycines, S. litura*, and *H. glycines*, respectively (Figure 2A). Interestingly, 57% (4/7) of the candidates against *S. litura* were also candidates against *A. glycines*. However, none of the candidate resistance genes against insects were shared with *H. glycines*, revealing a high intraclass (*i.e*., among insects), but no interclass overlap (*i.e*., among insects and nematodes). The shared genes are *Glyma.07G034400, Glyma.12G059900, Glyma.07G033100, Glyma.07G036400*, whose protein products are associated with phytohormone metabolism (KMD protein, Kelch repeat), transport (glucose and ATP transporters), and signaling (phospholipid:diacylglycerol acyltransferase), respectively. We also analyzed the overlap of pest resistance-related candidates with fungi resistance-related candidates from (Almeida-Silva and Venancio, 2021b) and found that a small number (n ≤5) of candidates against *H. glycines* and *A. glycines* are shared with *Fusarium* species (Figure 2B).

**Figure 2.**
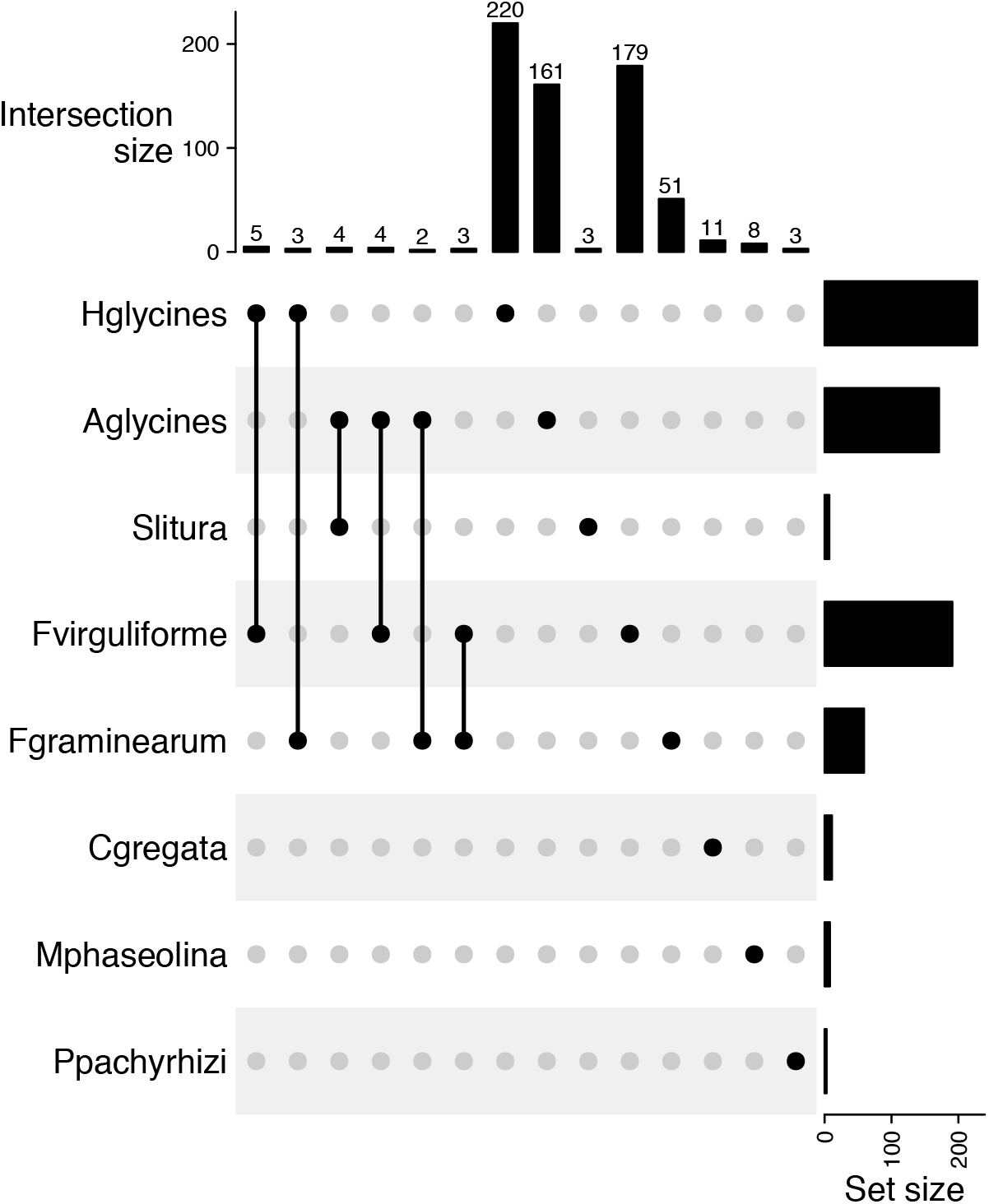
Upset plot with overlaps of candidate gene sets across fungal and pest species. Most candidate genes against *S. litura* are shared with *A. glycines*, suggesting a core defense against insects. However, insect resistance-related genes are not shared with nematode resistance-related genes. This suggests that insects and nematodes trigger different players of plant immunity. A small number of candidate resistance genes against pests are shared with *Fusarium sp*. resistance-related gene sets. Candidate resistance genes against fungi were retrieved from (Almeida-Silva and Venancio, 2021b).

The observed overlap of candidate gene sets for different insect species is desirable, because it suggests that shared candidates can be used in biotechnological applications to equip soybean accessions with broad-spectrum resistance (BSR) against insects. In our recent study on candidate resistance genes against fungi, we reported a highly species-specific response (Almeida-Silva and Venancio, 2021b), which seems to be a trend for filamentous pathogens (Ning and Wang, 2018; Kourelis and Van Der Hoorn, 2018). Conversely, BSR to insects has been reported more often and can be achieved with genes associated with the synthesis of volatile organic compounds and secondary metabolites, for instance (Dixit et al., 2013; Vosman et al., 2018). Altogether, these findings suggest that achieving BSR is a feasible approach to control pests in soybean fields.

### 3.3 Signaling, oxidative stress and transcriptional regulation shape soybean resistance to pests

We manually curated the high-confidence candidate resistance genes to predict the putative role of their products in plant immunity (Supplementary Table S4). Most of the prioritized candidates encode proteins involved in immune signaling (23%), oxidative stress (21%), and transcriptional regulation (16%) (Figure 3). Other candidates encode proteins that play a role in transport, translational regulation, physical defense, phytohormone and secondary metabolism, apoptosis, recognition, and direct function against pests (Figure 3).

**Figure 3.**
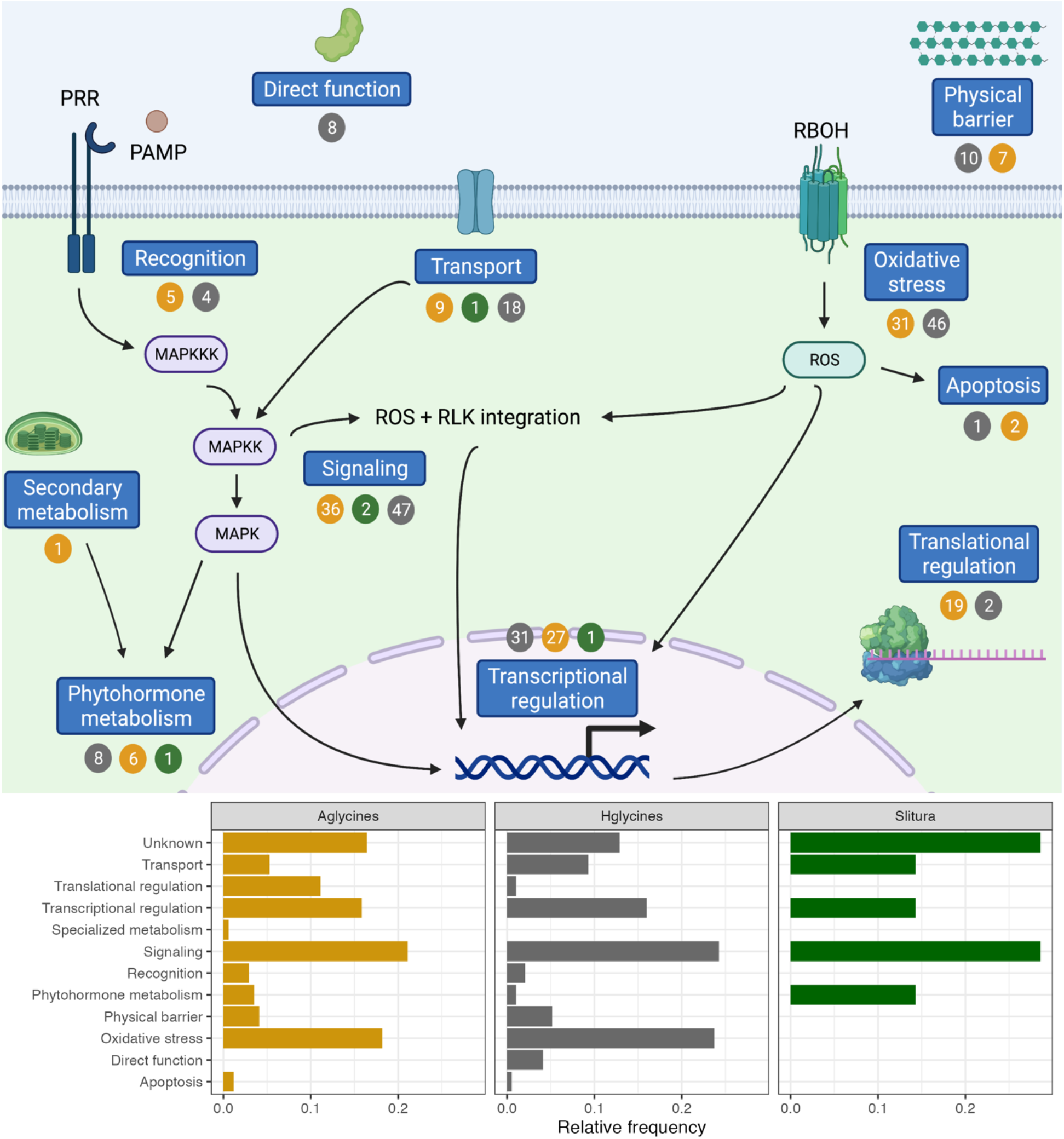
Prioritized candidate resistance genes and their putative role in plant immunity. Numbers in circles represent absolute frequencies of resistance genes against *Aphis glycines* (gold), *Heterodera glycines* (gray), and *Spodoptera litura* (green). PRR, pattern recognition receptor. PAMP, pathogen-associated molecular pattern. MAPKKK, mitogen-activated protein kinase kinase kinase. MAPKK, mitogen-activated protein kinase kinase. MAPK, mitogen-activated protein kinase. SAR, systemic acquired resistance. RBOH, respiratory burst oxidase homolog. ROS, reactive oxygen species. RLK, receptor-like kinase. Figure designed with Biorender (biorender.com).

Interestingly, 55 (15%) candidate genes lack functional description and, hence, we could not infer their roles in resistance (*n=*28, 25, and 2 for *A. glycines, H. glycines*, and *S. litura*, respectively). This finding demonstrates that our algorithm can also serve as a network-based approach to predict functions of unannotated genes, similarly to previous approaches (Almeida-Silva et al., 2020; Depuydt and Vandepoele, 2021). Genes encoding proteins of unknown function were in the top 4 most abundant categories for all species, revealing a potentially rich source of targets for biotechnological applications that would not have been identified if traditional SNP-to-gene mapping approaches were used.

We also developed a scheme that was used to rank high-confidence candidate genes (Table 2). As there are several candidate resistance genes against *A. glycines* and *H. glycines*, ranking candidates can help prioritize genes for experimental validation. Here, we suggest using the top 10 candidate resistance genes against each pathogen for experimental validation in future studies, which will likely reveal the most suitable candidates to develop soybean lines with increased resistance to each pest.

**Table 2.** Top 10 candidate resistance genes against each pest species and their putative roles in plant immunity. The predicted function for each gene was manually curated from the description of the best ortholog in *Arabidopsis thaliana*, using functional annotations from Soybase and TAIR.

| Gene | Predicted function | Resistance to | Role |
| --- | --- | --- | --- |
| Glyma.07G021800 | Chaperone, DnaJ domain | *A. glycines* | Translational regulation |
| Glyma.19G187200 | UDP-glycosyltransferase | *A. glycines* | Signaling |
| Glyma.10G002200 | Calmodulin | *A. glycines* | Signaling |
| Glyma.11G194500 | Acyl-CoA synthetase | *A. glycines* | Phytohormone metabolism |
| Glyma.01G210200 | Autophagy protein | *A. glycines* | Apoptosis |
| Glyma.04G085500 | Lysophospholipase | *A. glycines* | Signaling |
| Glyma.01G238800 | Sugar transporter | *A. glycines* | Transport |
| Glyma.19G018600 | AAA-ATPase | *A. glycines* | Oxidative stress |
| Glyma.13G326800 | Galactose oxidase | *A. glycines* | Physical barrier |
| Glyma.04G085600 | Unknown orphan gene | *A. glycines* | Unknown |
| Glyma.06G195300 | Protein of unknown function | *S. litura* | Unknown |
| Glyma.07G033100 | ATP transporter | *S. litura* | Transport |
| Glyma.07G034400 | KMD protein - Kelch repeat | *S. litura* | Phytohormone metabolism |
| Glyma.06G175400 | RNA-binding protein | *S. litura* | Transcriptional regulation |
| Glyma.05G194700 | Protein of unknown function | *S. litura* | Unknown |
| Glyma.07G036400 | Phospholipid:diacylglycerol acyltransferase | *S. litura* | Signaling |
| Glyma.12G059900 | Glucose transporter | *S. litura* | Transport |
| Glyma.18G077500 | Y-box transcription factor | *H. glycines* | Transcriptional regulation |
| Glyma.19G122700 | MYB transcription factor | *H. glycines* | Transcriptional regulation |
| Glyma.13G169900 | HD-Zip transcription factor | *H. glycines* | Transcriptional regulation |
| Glyma.09G245300 | MYB transcription factor | *H. glycines* | Transcriptional regulation |
| Glyma.01G179900 | Homeobox transcription factor | *H. glycines* | Transcriptional regulation |
| Glyma.16G053900 | TCP transcription factor | *H. glycines* | Transcriptional regulation |
| Glyma.01G207300 | HD-Zip transcription factor | *H. glycines* | Transcriptional regulation |
| Glyma.10G081700 | HOX transcription factor | *H. glycines* | Transcriptional regulation |
| Glyma.16G051800 | NAC transcription factor | *H. glycines* | Transcriptional regulation |
| Glyma.09G060200 | bHLH transcription factor | *H. glycines* | Transcriptional regulation |

### 3.4 Pangenome presence/absence variation analysis demonstrates that most prioritized genes are core genes

We analyzed PAV patterns for our prioritized candidate genes in the recently published pangenome of cultivated soybeans to unveil which soybean genotypes contain prioritized candidate genes and explore gene presence/absence variation patterns across genomes (Torkamaneh et al., 2021). We found that most candidates (98%) are present in all 204 accessions (Supplementary Figure 1A), similarly to what we observed for fungi resistance-related genes (Almeida-Silva and Venancio, 2021b). This unsurprising trend is likely due to the high level of gene content conservation in this pangenome, which has 91% of the genes shared by >99% of the genomes.

Further, we investigated if gene PAV patterns could be explained by the geographical origins of the accessions (Supplementary Figure 1B). As we observed in our previous study (Almeida-Silva and Venancio, 2021b), PAV patterns did not cluster by geographical origin, suggesting that gene PAV is not affected by population structure. As this pangenome comprises improved soybean accessions (Torkamaneh et al., 2021), the lack of population structure effect can be due to breeding programs targeting optimal adaptation to different environmental conditions (*e.g*., latitude and climate), even if they are in the same country.

### 3.5 Screening of the USDA germplasm reveals a room for genetic improvement

We inspected the USDA germplasm to find the top 10 most resistant genotypes against each pest species (see Materials and Methods for details). Strikingly, the most resistant genotypes do not contain all resistance alleles, revealing that, theoretically, they could be further improved to increase resistance (Table 3). None of the reported SNPs for resistance against *S. litura* have been identified in the SoySNP50k collection. Hence, we could not predict the most resistant accessions to this pest species.

**Table 3.** Top 10 most resistant soybean accessions against each pest species. Overall, the best genotypes do not reach the maximum potential. None of the resistance SNPs for *S. litura* have been identified in the USDA SoySNP50k compendium and, hence, we could not predict resistance potential against this species.

| Accession | Score | Potential | Species |
| --- | --- | --- | --- |
| PI468399A | 57 | 0.582 | *A. glycines* |
| PI532451 | 57 | 0.582 | *A. glycines* |
| PI468916 | 56 | 0.571 | *A. glycines* |
| PI468918 | 56 | 0.571 | *A. glycines* |
| PI479750 | 56 | 0.571 | *A. glycines* |
| PI479752 | 56 | 0.571 | *A. glycines* |
| PI468400B | 55 | 0.561 | *A. glycines* |
| PI468399B | 54 | 0.551 | *A. glycines* |
| PI507793 | 54 | 0.551 | *A. glycines* |
| PI479749 | 54 | 0.551 | *A. glycines* |
| PI556949 | 66 | 0.673 | *H. glycines* |
| PI84751 | 66 | 0.673 | *H. glycines* |
| Peking | 66 | 0.673 | *H. glycines* |
| PI438497 | 66 | 0.673 | *H. glycines* |
| PI548402 | 66 | 0.673 | *H. glycines* |
| PI438342 | 64 | 0.653 | *H. glycines* |
| PI549047 | 64 | 0.653 | *H. glycines* |
| PI597461C | 64 | 0.653 | *H. glycines* |
| PI404166 | 64 | 0.653 | *H. glycines* |
| PI437679 | 64 | 0.653 | *H. glycines* |

Our findings are in line with what we observed for resistance to fungi in the USDA germplasm (Almeida-Silva and Venancio, 2021b). Importantly, insect resistance potentials were lower than fungi resistance potentials (Wilcoxon rank-sum test, *P =* 7.7e-04), suggesting that pest resistance could be further improved. A feasible approach to increase pest resistance involves pyramiding quantitative trait loci (QTL) that confer partial resistance to each pest (Li et al., 2020). To accomplish this, the most resistant genotypes identified here can be used in breeding programs using marker-assisted selection or inspire CRISPR/Cas editing strategies to introduce beneficial alleles, leading to increased resistance. It is important to mention, however, that our model does not account for epistasis and different effect sizes of each variant. Hence, there might be accessions with a smaller number of SNPs with large effects that are more resistant than accessions with a greater number of SNPs with moderate effects.

### 3.6 Development of a user-friendly web application for network exploration

To facilitate network exploration and data reuse, we developed a user-friendly web application named SoyPestGCN (https://soypestgcn.venanciogroup.uenf.br/). Users can choose either the insect or the nematode GCN and input a soybean gene of interest (Wm82.a2.v1 assembly) to visualize the gene’s module, scaled intramodular degree, and hub status (Figure 4A). Additionally, users can explore enriched GO terms, Mapman bins and/or Interpro domains associated with the input gene’s module (Figure 4). This resource can be particularly useful for researchers studying soybean response to other pest species, as they can check if their genes of interest are located in defense-related coexpression modules. Also, researchers studying other species can verify if the soybean ortholog of their genes of interest is located in a defense-related module. The application is also available as an R package named SoyPestGCN (https://github.com/almeidasilvaf/SoyPestGCN). This package lets users run the application locally as a Shiny app, ensuring the application will always be available, even in case of server downtime.

**Figure 4.**
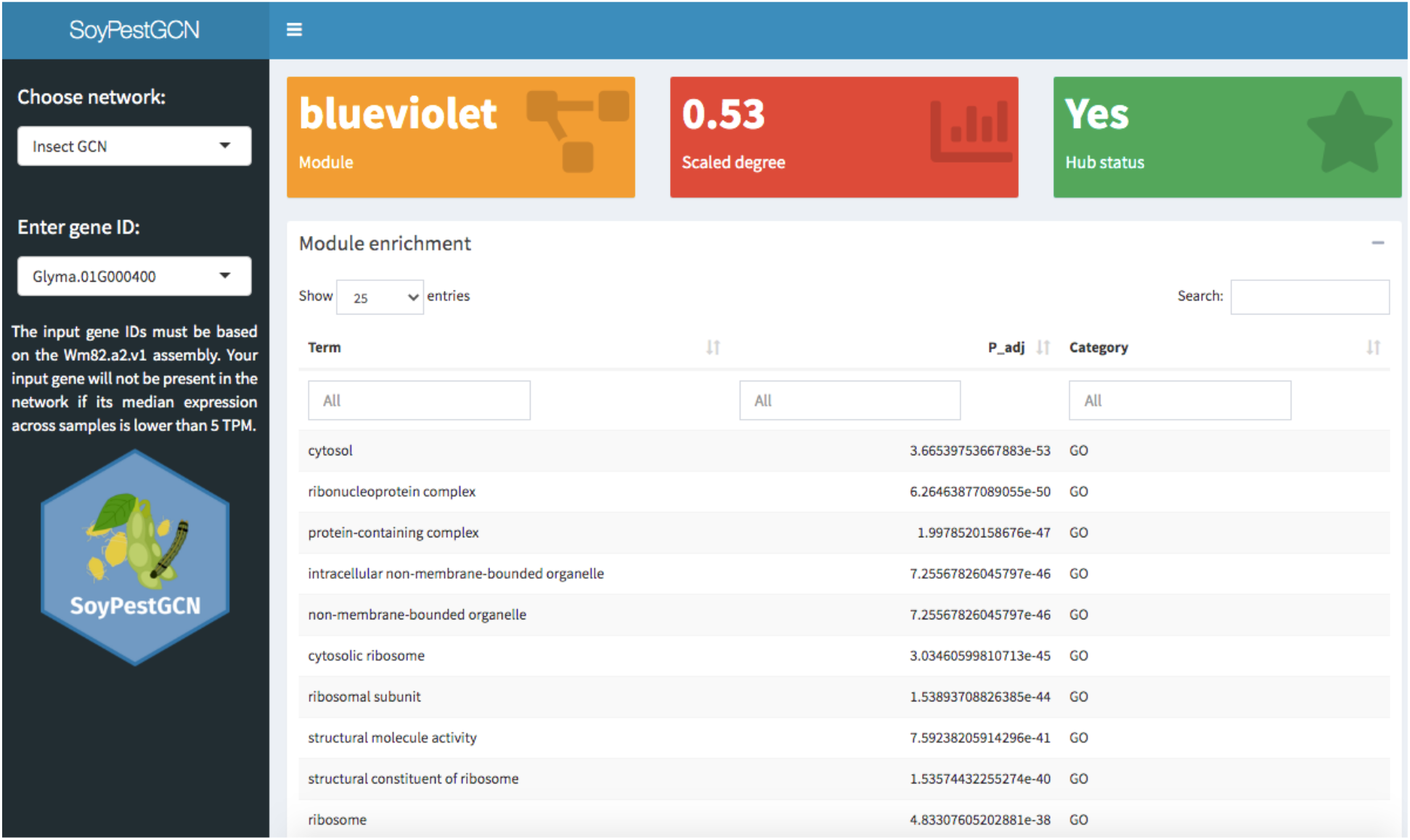
Functionalities in the SoyPestGCN web application. Screenshot of the page users see when they access the application. In the sidebar, users can specify either the insect GCN or the nematode GCN followed by the ID of a gene of interest (Wm82.a2.v1 assembly). For each gene, users can see the gene’s module (orange box), scaled degree (red box), hub gene status (green box), and an interactive table with enrichment results for MapMan bins, Interpro domains and Gene Ontology terms associated the gene’s module. P-values from enrichment results are adjusted for multiple testing with Benjamini-Hochberg correction.

## 4 Conclusions

By integrating publicly available GWAS and RNA-seq data, we found promising candidate genes in soybean associated with resistance to three pest species, namely *A. glycines, S. litura*, and *H. glycines*. The prioritized candidates encode proteins involved in immunity-related processes such as in recognition, signaling, transcriptional regulation, oxidative stress, specialized metabolism, and physical defense. We have also found the top 10 most resistant soybean accessions against each pest species and hypothesize that they can be used in soybean improvement programs, either via breeding with marker-assisted selection or through genome editing. The coexpression network generated here was also made available in a web resource and R package to help in future studies on soybean-pest interactions.

## Data availability

All data and code used in this study are available in our GitHub repository (https://github.com/almeidasilvaf/SoyPestGCN_paper) to ensure full reproducibility.

## Acknowledgements

This work was supported by Fundação Carlos Chagas Filho de Amparo à Pesquisa do Estado do Rio de Janeiro (FAPERJ; grants E-26/203.309/2016 and E-26/203.014/2018), Coordenação de Aperfeiçoamento de Pessoal de Nível Superior - Brasil (CAPES; Finance Code 001), and Conselho Nacional de Desenvolvimento Científico e Tecnológico. The funding agencies had no role in the design of the study and collection, analysis, and interpretation of data and in writing.

## Author contributions

Conceived the study: FA-S and TMV. Data analysis: FA-S. Funding, project coordination and infrastructure: TMV. Manuscript writing: FA-S and TMV.

## Conflicts of interest

none.

## Notes

https://soypestgcn.venanciogroup.uenf.br/

### Competing Interest Statement

The authors have declared no competing interest.

